# Multiplex recombinase polymerase amplification (RPA) assay developed using unique genomic regions and coupled with a lateral flow device for rapid on-site detection of genus *Clavibacter* and *C. nebraskensis*

**DOI:** 10.1101/2020.08.22.262824

**Authors:** Adriana Larrea-Sarmiento, James P. Stack, Anne M. Alvarez, Mohammad Arif

**Affiliations:** Department of Plant and Environmental Protection Sciences, University of Hawaii at Manoa, Honolulu, Hawaii - 96822, United States of America; Department of Plant Pathology, Kansas State University, Manhattan, Kansas – 66506, United States of America

**Keywords:** Bacteria, *Clavibacter*, diagnosis, plant disease, multiplex RPA

## Abstract

*Clavibacter* is an agriculturally important bacterial genus comprising nine host-specific species/subspecies including *C. nebraskensis* (*Cn*), which causes Goss’s wilt and blight of maize. A robust, simple, and field-deployable method is required to specifically detect *Cn* in infected plants and distinguish it from other *Clavibacter* species for quarantine purposes and timely disease management. A multiplex Recombinase Polymerase Amplification (RPA) coupled with a Lateral Flow Device (LFD) was developed for sensitive and rapid detection of *Clavibacter* and *Cn* directly from infected host. Unique and conserved genomic regions, the ABC transporter ATP-binding protein CDS/ABC-transporter permease and the MFS transporter gene, were used to design primers/probes for specific detection of genus *Clavibacter* and *Cn*, respectively. The assay was evaluated using 52 strains, representing all nine species/subspecies of *Clavibacter*, other closely related bacterial species, and naturally- and artificially-infected plant samples; no false positives or negatives were detected. The RPA reactions were also incubated in a closed hand at body temperature; results were again specific. The assay does not require DNA isolation and can be directly performed using host sap. The detection limit of 10 pg and 100 fg was determined for *Clavibacter*- and *Cn*-specific primers/probes, respectively. The detection limit for *Cn*-specific primer/probe set was decreased to 1,000 fg when 1 µL of host sap was added into the RPA reaction containing 10-fold serially diluted genomic DNA; though no effect was observed on *Clavibacter*-specific primer/probe set. The assay is accurate and has applications at point-of-need diagnostics. This is the first multiplex RPA for any plant pathogen.

**IMPORTANCE:** *Clavibacter* species are prevalent worldwide as have the potential to result in systemic infection. In the past, detection attempts have relied on both molecular- and immunological-based assays; however, current detection methods are time consuming and laborious. Field-deployable tests are desirable to identify potential samples infected with *Clavibacter* species. This study demonstrates that the field-deployable isothermal multi-target recombinase polymerase amplification can be performed for the simultaneous detection of the genus *Clavibacter* in general (all species), and *C. nebraskensis*, in particular, without specialized equipment. Additionally, the multiplex RPA coupled with a LFD may confer the benefits of faster detection and discrimination of *Clavibacter* species that affect critical regions susceptible to infection. This user-friendly format offers a flexible assay to complement both nucleic acid amplification and novel diagnosis methods without the need for DNA purification; this assay may serve as a point-of-reference for developing multiplex RPA assay for other plant pathogens.

## INTRODUCTION

*Clavibacter* genus is a well-known, Gram-positive plant-pathogenic bacterium belonging to the *Microbacteriaceae* family—contains high GC—and is responsible for several devastating diseases in staple crops worldwide (1, 2). Previously, this genus consisted of only one species subdivided into nine subspecies, but recently, six subspecies were elevated to the species level (3). The genus now includes *C. michiganensis* (*Cm*), *C. sepedonicus* (*Cs*), *C. insidiosus* (*Ci*), *C. tessellarius* (*Ct*), *C. capsici* (*Cc*), *C. nebraskensis* (*Cn*), affecting tomato, potato, alfalfa, wheat, peppers, and corn, respectively. However, the other three subspecies—*C. michiganensis* ssp. *chiloensi*s (*Cmch*) and *C. michiganensis* ssp. *californiensis* (*Cmca*) isolated from tomato, and *C. michiganensis* ssp. *phaseoli* (*Cmp*) from beans—have characteristics that warrant their elevation to the species level as well (M. Arif; unpublished information).

All six species and three subspecies are host specific, infecting the main host and a few closely related species. *Clavibacter* species and subspecies are usually disseminated through contaminated seed, propagative materials and soil; secondary infections usually occur through wounds but in some cases stomates or hydathodes. In the early stages of the infection, all members within the *Clavibacter* genus reside as biotrophs within xylem vessels eventually leading to systemic disease (1, 4). Three main factors, including the severity of the disease, the pathogen’s ability to invade seeds, and latent symptomless infections, have necessitated the classification of *Cm, Cs*, and *Ci* as quarantine organisms under the European Union Plant Health Legislation, as well as in many other countries (2). Despite numerous attempts to breed resistant varieties, no resistant plant cultivars have been commercialized. Due to adverse effects of *Clavibacter* species on economically important crops, a field-deployable, cost-effective, sensitive and specific detection tool is needed to detect early infections by *Clavibacter* species at the point-of-need (PON, 1, 5-8).

Although suitable multiplex molecular techniques are available to detect multiple target pathogens, these techniques are restricted to centralized laboratories and suffer from several other limitations (9-11). To address the demand of point-of-need and integrated field-applicable technologies, isothermal nucleic acid amplification methods have been the object of extensive research efforts; many isothermal methods are available, but among them, loop-mediated isothermal amplification (LAMP;12) and recombinase polymerase amplification (RPA;13) are the most commonly used techniques (2, 14-20). RPA effectiveness is highlighted by rapid detection of multiple targets within a single reaction; however, additional advantages of this technique include the availability of lyophilized reagents, obviation of initial denaturation, high sensitivity, affordability, reduced equipment requirements, and operation at constant low temperature employing recombinase-primer complexes (11, 14, 16, 20-23).

RPA possesses an inherent tolerance for sample impurities that may inhibit other platforms based on nucleic acid amplification. Additionally, easy visualization of the results using lateral flow devices can be attained without the involvement of fluorescence readers (16, 20), which demonstrates the flexibility of RPA for rapid pathogen diagnosis. RPA detection can be achieved as early as in ten minutes, and measurements can be obtained over an extended range of temperatures with a detection limit of 1 fg (10, 16, 20, 24-25). It has been reported that even greater optimization of this method can be achieved to prevent false negatives and augment the sensitivity on the multiplex format, although such effects are contingent upon the characteristics of the target sequences, amplicon sizes, primer, and probe design ratios (11, 24, 26-27). While RPA assays are widely used in the detection of animal and human pathogens, its use in plant pathogen detection is limited, but increasing with the availability of commercial kits (14, 16, 20, 28-32); yet, no RPA multiplex has been developed for any plant pathogens.

To address the demand for a point-of-need test, we proposed the development of an isothermal multiplex recombinase polymerase amplification assay, not only noted for a high degree of sensitivity and specificity with the potential to be implemented in a laboratory setting, but also for the portable format that enables on-site detection and differentiation of *Clavibacter* species from *C. nebraskensis*. This protocol will also serve as a point-of-reference to develop multiplex RPA assays, coupled with a lateral flow device, for other plant pathogens.

## MATERIALS AND METHODS

### Bacterial Isolates and DNA Extraction

Strains were obtained from the Pacific Bacterial Collection at the University of Hawai’i at Manoa. Thirty-two bacterial strains representing all species/subspecies within the genus *Clavibacter* were used in the inclusivity panel. The exclusivity panel was comprised of twenty strains, including Gram-positive and Gram-negative bacterial strains, and characterized as either related or disparate plant pathogens affecting the same crops as the *Clavibacter* spp. (Table 1). Almost the same strain panels were used by Larrea-Sarmiento et al (2019; 33) to validate a multiplex qPCR assay developed for *Clavibacter* species and *C. nebraskensis.*

**Table 1.**
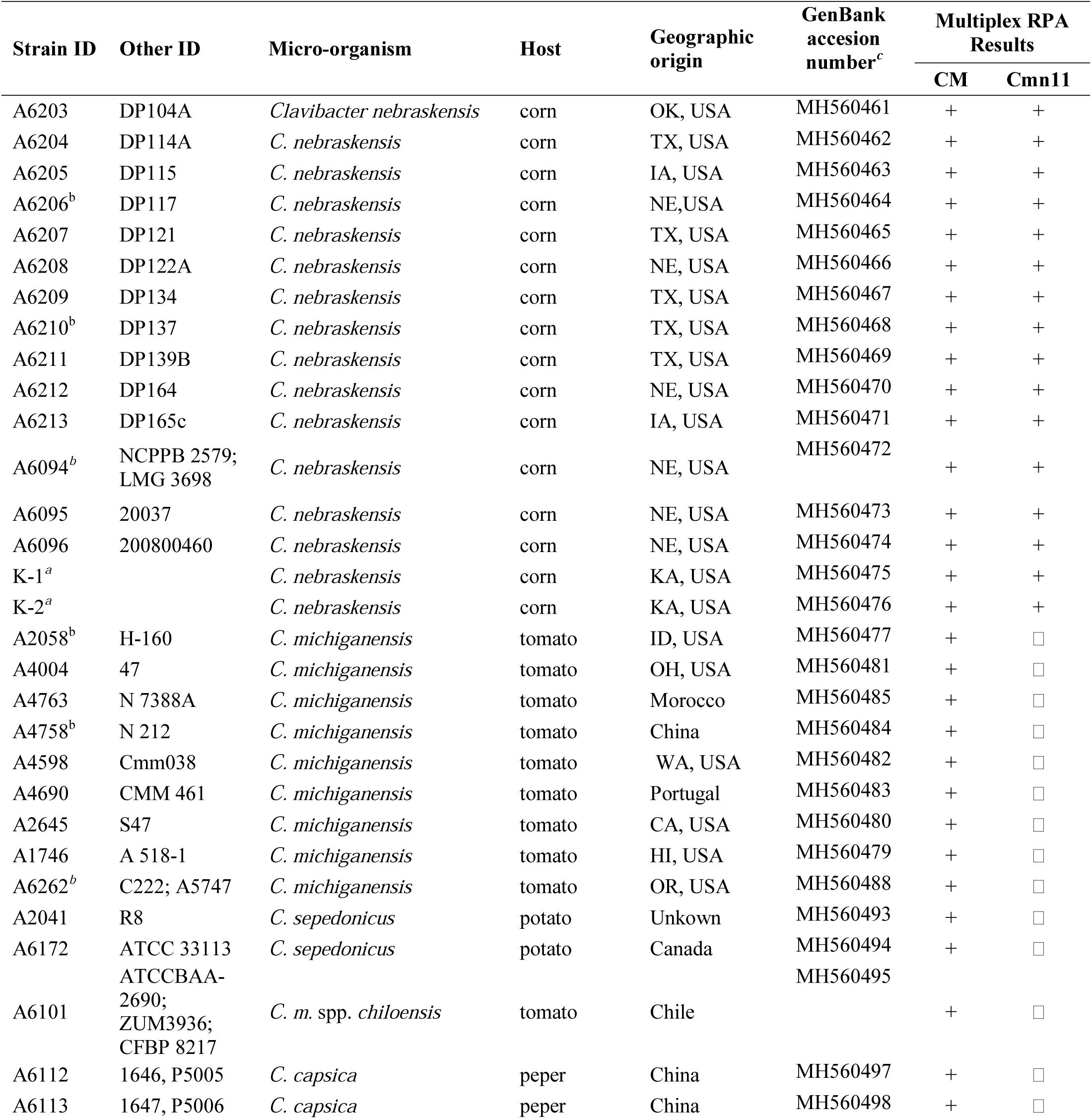
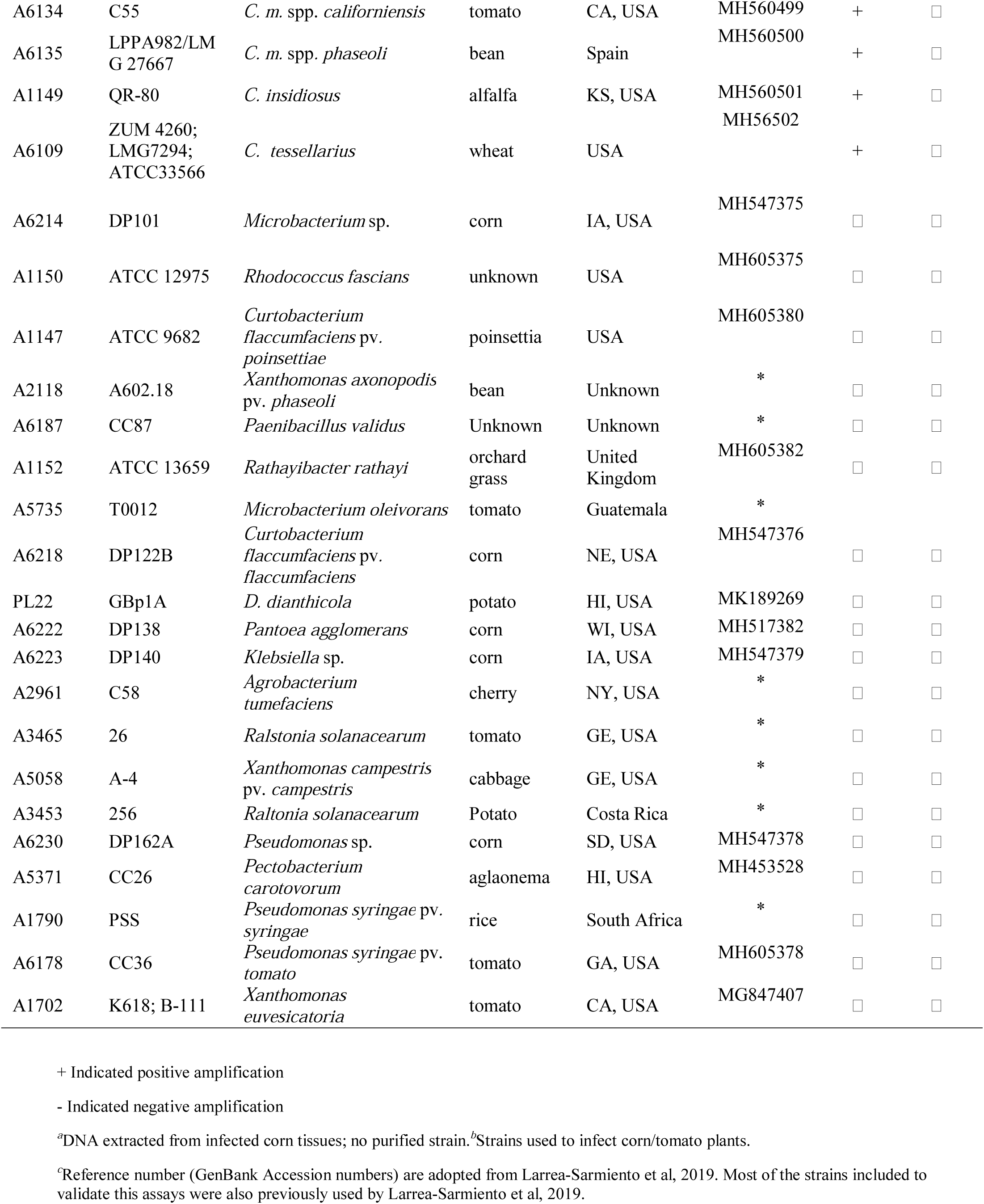
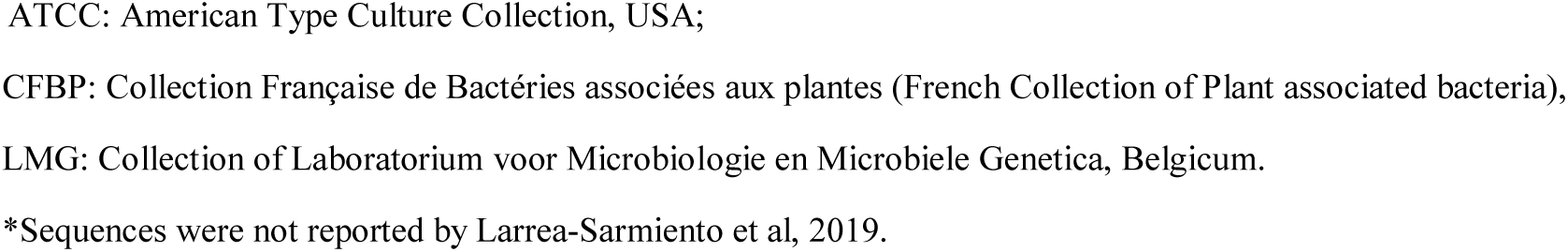
Details of the bacterial strains used in inclusivity and exclusivity panels to validate the multiplex RPA assay for specific detection of genus *Clavibacter* in general and *C. nebraskensis*.

Gram-positive and gram-negative strains were cultured on TZC-S and TZC media, respectively (34). Plates were incubated for 2 days at 26°C (±2°C) for DNA extraction or colony PCR. Bacterial genomic DNA was extracted using DNeasy UltraClean Microbial Kit (MO BIO; QIAGEN, Germantown, MD) according to the manufacturer’s instructions. DNA was quantified using the NanoDrop v.2000 spectrophotometer (Thermo Fisher Scientific, Inc., Worcester, MA) for further sensitivity assays. DNA of naturally infected plant samples were received from Kansas State University (Jarred Yasuhara-Bell and James P. Stack;33).

### Target Selection and Primer and Probe Design

Unique genomic regions were identified using the comparative genomic analyses previously reported (33); ABC transporter ATP-binding protein CDS/ABC-transporter permease and MFS transporter were used for genus *Clavibacter* and *C. nebraskensis*, respectively. Primers and probes were designed following the instructions provided by TwistDx Ltd. (Maidenhead, UK), with lengths of 32 bp and 49 bp, respectively. Primers and probes specificity was evaluated *in silico* using the NCBI GenBank BLASTn tool (35). The primers and probes were also aligned with *Clavibacter* species genomes using Geneious version 7.1. Details of primers and probes, synthesized by Integrated DNA Technologies (IDT, Coralville, IA) and LGC Biosearch Technologies (Petaluma, CA), respectively, are provided in Table 2. Reverse primers were labeled with fluorescent 6-FAM (6-Carboxyfluorescein) and DIG (Digoxigenin) while both probes were labeled with biotin.

**Table 2.**
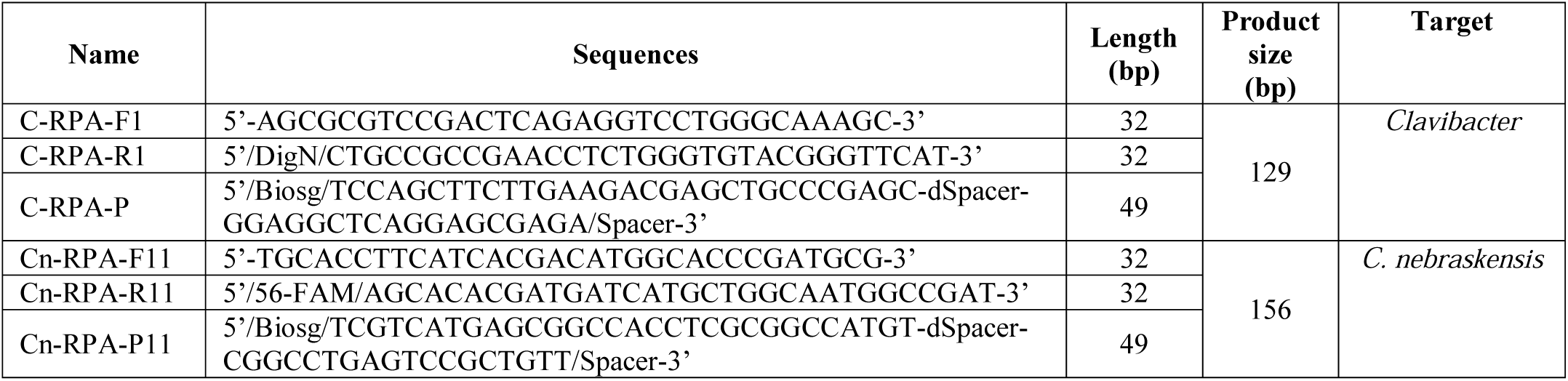
Details of multiplex RPA primers and probes developed for specific detection of *Clavibacter* in general and *C. nebraskensis*

Efficiency and specificity of the primers were first evaluated in single end-point PCR reactions with a final volume of 20 µL, containing 1 µL of both sense and anti-sense primers (5 mM), 10 µL of GoTaq Green Master Mix, 7 µL nuclease free-water, and 1 µL of genomic DNA. PCR assays were performed with the following conditions: initial denaturation at 95 °C for 5 min, with 35 cycles of 94 °C for 45 secs, 60 °C for 1 min and 72 °C for 1 min, and a final extension at 72 °C for 5 min.

### Single RPA Assay Design

TwistAmp nfo kits were supplied by TwistDx Ltd., Cambridge, United Kingdom. First, single RPA reactions were performed to validate the specificity of each primer/probe set. The components and conditions were followed as recommended by the manufacturer with slight modifications: each reaction containing an RPA TwistAmp nfo freeze-dried enzyme pellet was rehydrated with 45.5 µl of master-mix consisting of 29.5 µl rehydration buffer, 2.1 µl of each reverse/forward primer set (10 µM), 1.2 µl of probe (5 µM) and 10.6 of nuclease-free water. To each reaction tube, 2 µl of either purified DNA or crude plant extract was added followed by 2.5 µl of Mg-acetate; the tube was mixed by inversion and brief centrifugation.

### Multiplex RPA Assay Optimization

The multiplex RPA assays were optimized in three phases using the TwistAmp nfo kit. For initial optimization, RPA was run at two incubation temperatures, 37 °C and 39 °C, and amplicons resolved in 1.5% agarose gels to identify the optimal amplification temperature.

Upon selection of the optimal temperature, primers (10 mM)/probe (5 µM) concentrations were evaluated in three ratios in multitarget assays 1-low (*Cn* 1.2 µL/1.4 µL; *Clavibacter* 0.8 µL /0.6 µL), 2-equal (*Cn* 1.2 µL/0.8 µL; *Clavibacter* 1.2 µL/0.8 µL) and 3-high (*Cn* 1.5 µL/0.9 µL; *Clavibacter* 1.0 µL/0.6 µL). The third phase of assay optimize determined optimum incubation time. Multiplex RPA assays were performed using pure DNA extracted from *C. nebraskensis* with incubation periods ranging from 5 to 30 min.

### Sensitivity and Spiked Multiplex RPA Assays

Two independent assays targeting *Clavibacter* and *Cn* were integrated into one multiplex assay. Multiplex RPA assay components were: rehydrating RPA Twist-nfo pellet with 45.5 µL of master mix containing 1.2 µL each RPA primer (total 4 primers were added, 10 µM), 0.8 µL each probe (5 µM; total two probes), 29.5 µL 1X rehydration buffer and 9.6 µL nuclease-free water. A volume of 2.5 µL of MgOAc applied to the lid, and 2 µL of DNA was added to the final reaction; the reaction was mixed and briefly centrifuged before incubation. For each RPA assay, the relative concentration of purified genomic DNA was measured using the NanoDrop v.2000 spectrophotometer. The limit of detection of RPA multiplex assays was determined using serially diluted genomic DNA from 10 ng to 100 fg (A4757-*C. michiganensis* and A6094-*C. nebraskensis*). Effect of plant extract was evaluated by performing a spiked test: one µL of template DNA was used from each 10-fold serial dilution of pure bacterial genomic DNA and mixed with 1 µl of crude extract from a healthy corn leaf tissue. Finally, 2 µL of the template was used from each spiked dilution in an RPA reaction with a total volume of 50 µL. Nuclease-free water and/or healthy corn tissue were included in every run as negative controls for the DNA and spiked sensitivity assays, respectively.

### Field-applicability

In order to test the specificity of RPA assay as a method for point-of-need detection, a total of six strains, three *Cm* A2058, A4758, A5747, and three *Cn* A6206, A6210, A6094, were selected for inoculation of tomato and corn plants, respectively. Strains were grown on YSC medium, and bacterial inoculum was prepared using PBS buffer pH 7 to a final concentration of 10^8^ cells per ml. For *Cn*, corn plants in the V3 developmental stage were inoculated at the third leaf as described by Ahmad et al. (2005; 36). For *Cm*, the stems of three-week-old tomato plants were pierced between the two cotyledons with a needle as described by Kaneshiro et al. (2006; 37); 10 µl of *Cm* inoculum was poured directed onto the wound. For both *Cn* and *Cm* assays, a negative control was mock-inoculated with 10 ul of PBS alone. Inoculated and control plants were placed into translucent plastic bags for 24 hours to maintain humidity. Plant tissues were collected after 30 days-post-inoculation (33).

A master mix containing all RPA reagents, except DNA template (2.0 µl) and MgOAc was prepared, as described above and distributed into individual RPA reaction tubes containing a dried enzyme pellet. Field applicability of multiplex RPA reaction was tested by grinding 100 mg each of healthy and infected corn leaf tissue, and healthy and infected tomato stem tissue in 200 µL of TE buffer; the crude sap/extract was used as a template without isolating the DNA. For each reaction, 2 µL of crude extract was added followed by 2.5 µL MgOAc. The contents were thoroughly mixed by shaking and immediately incubated in a closed hand for 30 minutes.

Portable Lateral flow PCRD Nucleic Acid Detectors containing two different detection lines and one positive control line were coupled to visualize the results. A total of 5 µL of RPA product was added to 70 µL of PCRD extraction buffer, and a total of 75 µl were applied to the receptacle of a portable lateral flow device. Carbon particles at the capture lines of the lateral flow device enabled specific detection of amplicons labeled with either DIG/Biotin and/or FAM/Biotin within 10 min, and the results were visible to the unaided eye.

## RESULTS

### *In silico* Specificity Validation

We developed two unique primer and probe sets for specific detection of the genus *Clavibacter* and *C. nebraskensi*s (Figure 1). The genomic regions, ABC transporter ATP-binding, sugar ABC transporter permease, and MFS transporter, were identified through comparative genomic analyses (33). Developed primers/probe sets showed 100% query coverage and 100% identity with only the target species; *Clavibacter* primer/probe regions were present in all species within the genus *Clavibacter*, while the *Cn* primer/probe region was exclusively present in *C. nebraskensis*. Initial specificity tests were performed with endpoint PCR and showed no false positives or negatives (Fig. S1).

**Figure 1.**
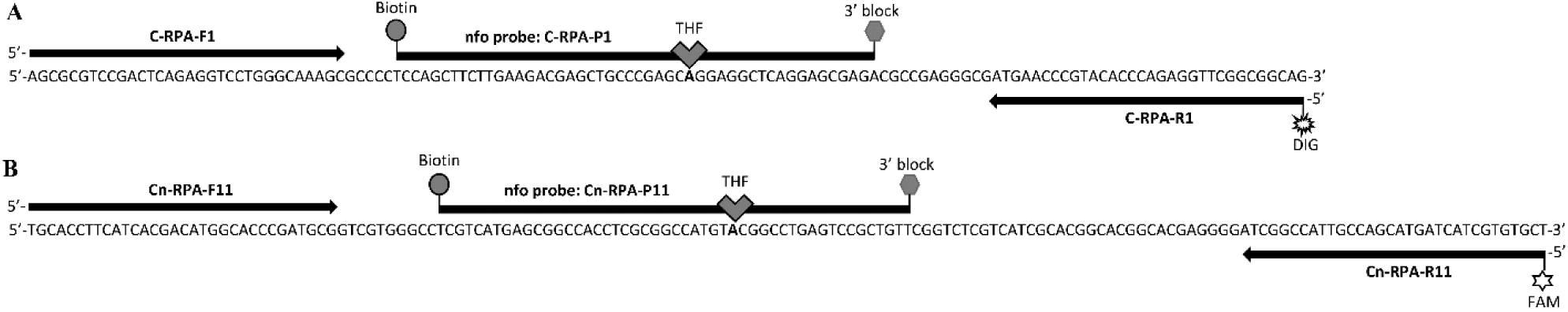
Multiplex RPA primer and probe locations developed to specifically detect genus *Clavibacter* (A) and *C. nebraskensis* (B). (A) Target fragment size 129 bp, reverse primer labeled with DIG; (B) Target fragment size 156 bp, reverse primer labeled with FAM. The probe and primer are 49- and 32-bp long, respectively. THF - tetrahydrofuran residue; DIG – digoxigenin; FAM - 6-carboxy-fluorescein.

### RPA Assay Optimization

The RPA reaction yielded greater amplification effects (thicker bands) when incubated at 39 °C compared to 37 °C (data not shown). Primer/probe set concentration was also optimized; the results indicated that equal concentrations of primers (1.25 µL, 10 mM) and probes (0.8 µL, 5 µM) gave the best results (Fig. S2). Finally, the effect of RPA incubation time was evaluated from 5 to 30 minutes. The results indicated that a positive amplification can be obtained in 10 min but with a low band intensity (Figure 2). However, the 20-30 min incubation time delivered optimum results (Figure 2). In addition, a brief spin after 4 minutes once the RPA reaction started showed to have better results compared with no spin.

**Figure 2.**
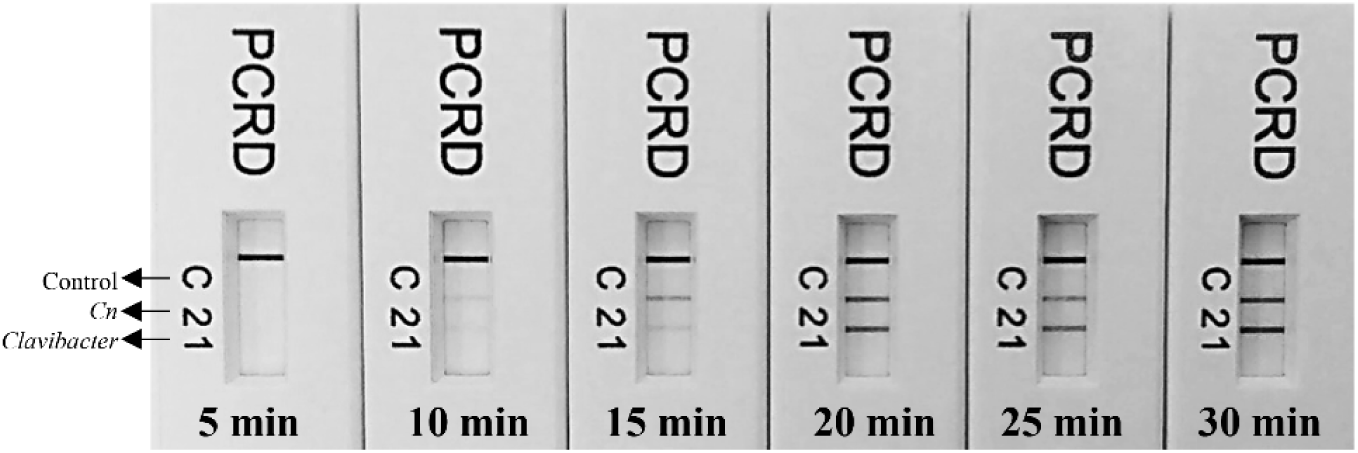
Determination of optimum time for successful detection of the genus *Clavibacter* and *C. nebraskensis* using the developed multiplex RPA assay. Line 1: detects *Clavibacter* species/subspecies strains; line 2 detects *C. nebraskensis* strains, and; line C is a control line. The primer/probe sets for genus *Clavibacter* and *C. nebraskensis* are labeled with Biotin/DIG and Biotin/FAM, respectively. The detection can be achieved within 10 min but the optimum line intensity on PCRD was observed in 20-30 min.

### RPA Assay Selectivity and Reaction Rate

RPA single and multiplex reactions demonstrated high levels of sensitivity and specificity. No false positives were reported using *Clavibacter-* or *Cn-*specific primer/probe sets (Figure 3). All six reported species including *Cm, Cs, Ci, Ct, Cc, Cn*, and three subspecies *Cmch, Cmca*, and *Cmp* within the genus *Clavibacter* were detected. *Clavibacter*-specific primer/probe labeled with DIG/Biotin was sufficiently robust to detect all species and subspecies within the genus *Clavibacter*, whereas, *Cn-*specific primer/probe labeled with FAM/Biotin showed specific detection of *C. nebraskensis* strains without displaying cross-reactions with other species/subspecies within the genus *Clavibacter*. Table 1 shows the results of inclusivity and exclusivity tests. All 32 strains representing all species/subspecies within genus *Clavibacter* were detected using the genus-specific primer/probe set; while a total of 16 samples (14 from pure culture and 2 from naturally infected tissues) were accurately identified as *C. nebraskensis* using *Cn*-specific primer/probe set. Results were visible to the unaided eye using the PCRD Nucleic Acid Detector. Lines 1, 2, and C correspond to *Clavibacter* species, *Cn*, and control line, respectively (Figure 3).

**Figure 3.**
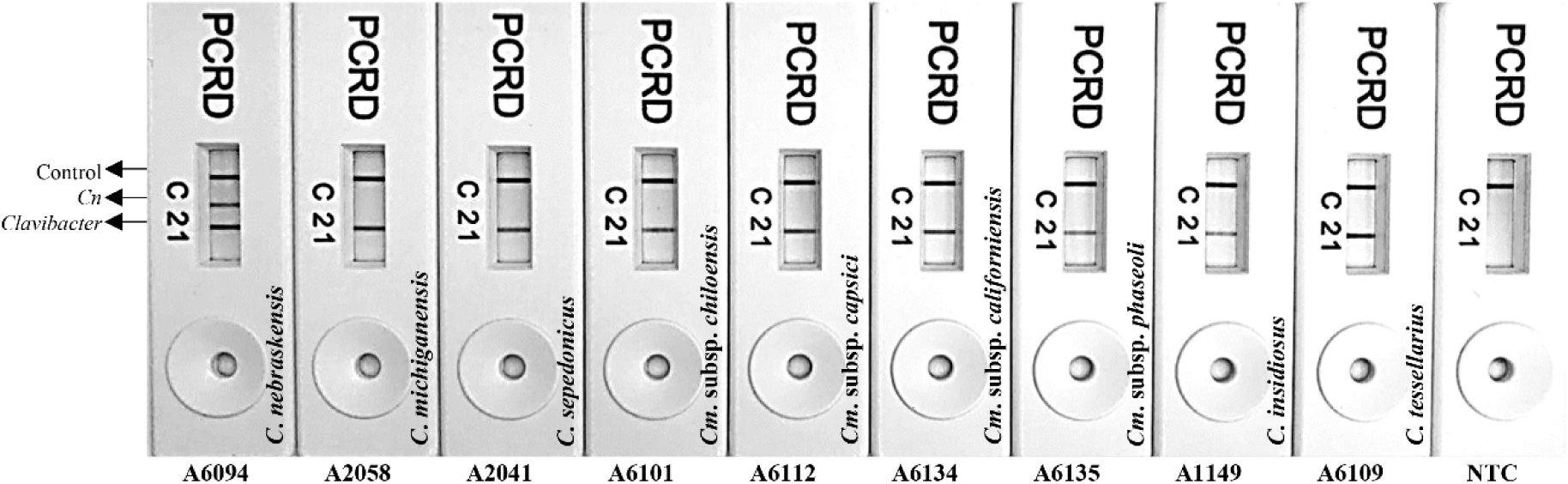
Multiplex RPA assay validation with all known species within the genus *Clavibacter*. PCRD Nucleic Acid Detector allows the detection of labeled RPA amplicons: line 1-detects the nine DIG/Biotin-labelled *Clavibacter* species/subspecies amplicons; line 2 -specific detection of FAM/Biotin-labeled *C. nebraskensis* amplicons, and; line C - control line. NTC (non-template control; water) is included in the last reaction. Strain numbers and species/subspecies names are provided in the figures.

### RPA Limit of Detection

Three analytical sensitivity assays were performed to assess the detection limit of the developed multiplex RPA assay (Figure 4). For each RPA assay, 2 µL of 10-fold serially diluted genomic DNA was added as a template. In the first experiment, the limit of detection was determined using the DNA extracted from pure *Cn* culture; a detection limit of 100 fg and 10 pg 10 pg and 100 fg was observed for *Clavibacter*- and *Cn*-specific primer/probe sets, respectively. In the second experiment, the inhibitory effect of plant inhibitors was evaluated, 1 µL of corn plant extract was added to each reaction, containing 10-fold serially diluted DNA and other RPA reaction components; a detection limit of 10 pg and 1 pg was observed for *Clavibacter*- and *Cn*-specific primer/probe sets, respectively. In a 3^rd^ assay, pure genomic DNA from *C. michiganensis* was serially diluted and used for sensitivity assay; detection limit of 10 pg was observed with *Clavibacter*-specific primer/probe set while, as expected, no amplification with *Cn*-specific primer/probe primer/probe set (Figure 4).

**Figure 4.**
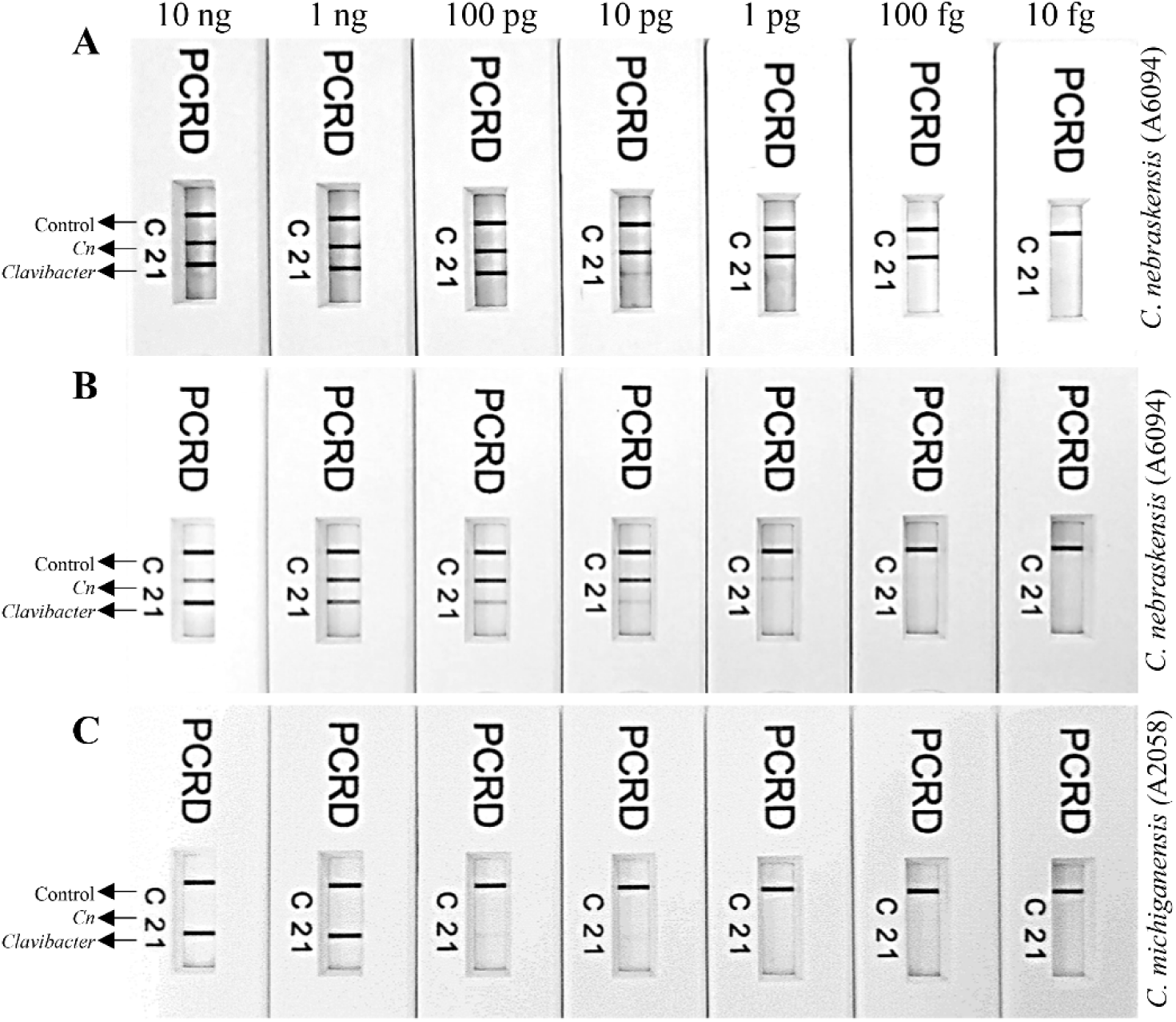
Limit of detection determination of multiplex RPA assays developed for specific detection of genus *Clavibacter* in general and *C. nebraskensis*. Line 1: detects *Clavibacter* species/subspecies strains; line 2 detects *C. nebraskensis* strains, and; line C is a control line. The primer/probe sets for genus *Clavibacter* and *C. nebraskensis* are labeled with Biotin/DIG and Biotin/FAM, respectively. A 10-fold genomic DNA serial dilution (10 ng – 10 fg) was used to perform the sensitivity assays. (**A)** Limit of detection determination using pure *C. nebraskensis* (A6094) DNA; detected until 100 fg and 10 pg using *Clavibacter* and *C. nebraskensis* specific primers/probe set, respectively. **(B)** Spiked multiplex test performed by adding 1 µL of corn plant sap in 10-fold serially diluted *C. nbraskensis* (A6094) genomic DNA; detected until 1,000 fg and 10 pg using *Clavibacter* and *C. nebraskensis* specific primers/probe set, respectively. **(C)** Detection using 10-fold serially diluted *C. michiganensis* (A2058) purified genomic DNA; detected until 10 pg by *Clavibacter* genus-specific primers/probe—no detection with *C. nebraskensis* specific primers/probe set.

### Detection From Artificially and Naturally Infected Plant Samples and Field Applicability

Determination of specificity and detection range of RPA, entailed for point-of-need assays, were carried out using tomato and corn tissues collected from symptomatic 30 dpi plants inoculated with *Cm* and *Cn*, respectively. Mock inoculated plant tissues served as a negative control. To evaluate the robustness of the developed RPA assay, no DNA purification was performed— crude sap from infected plants was used directly in the RPA reaction. Testing additional parameters for field viability, temperature conditions, and incubation time were evaluated, and RPA assays were performed in the aplm of a closed hand without employing additional equipment. Positive results were observed from tomato and corn tissues infected with *Cm* (only with *Clavibacter-*specific assay) and *Cn.* Presence of pathogens from infected plants was cross-confirmed using end-point PCR (data not shown). Neither positive results nor cross-amplification were observed when mock-inoculated tissues were tested on multiplex reactions (only the control line was observed) (Figure 5). These results confirmed that the developed RPA assay can specifically detect any species/subspecies within the genus *Clavibacter* directly from infected plant materials without DNA purification or specialized equipment. Additionally, portable Lateral flow PCD Nucleic Acid Detectors from Abingdon Health help to detect multiple targets without exhibiting false positive or false negative results, producing a robust multi-target RPA assay.

**Figure 5.**
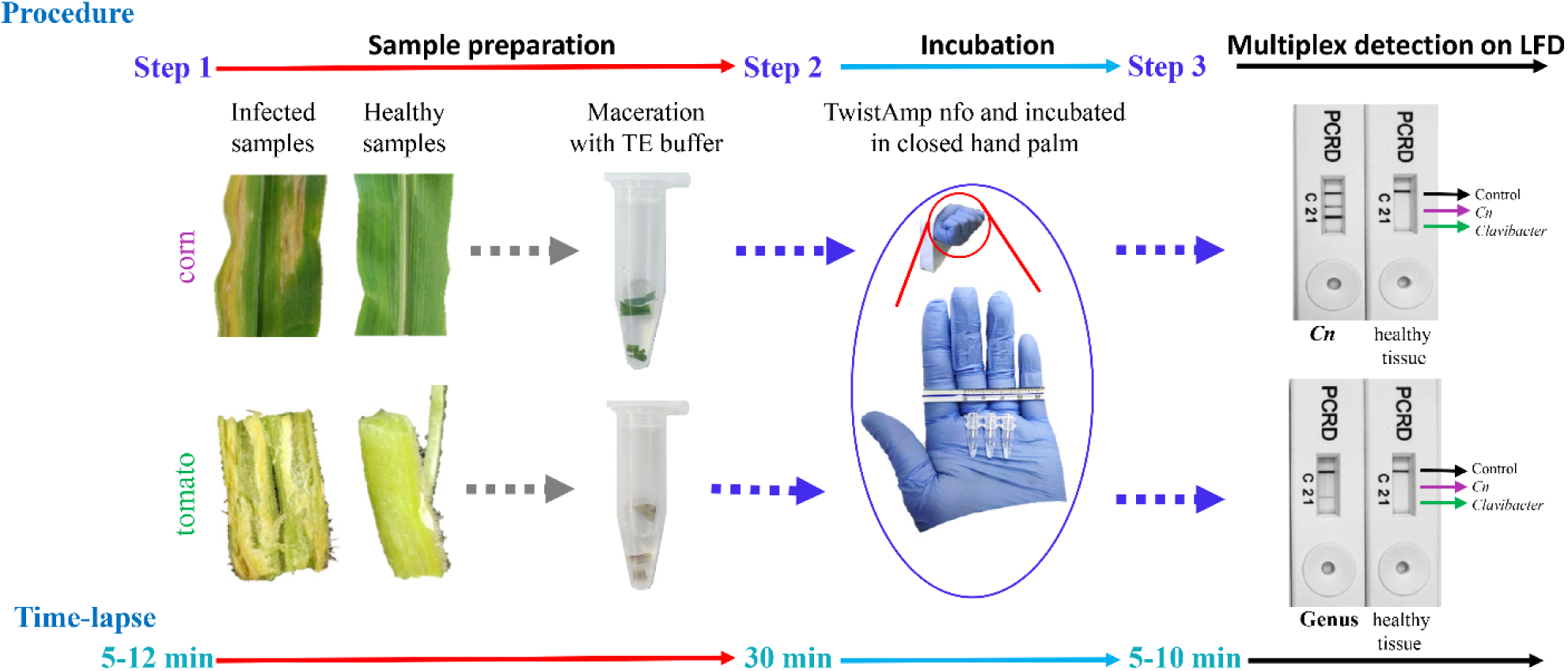
Schematic steps involve performing multiplex RPA intend for on-filed detection of genus *Clavibacter* in general and *C. nebraskensis*. Line 1: detects *Clavibacter* species/subspecies strains; line 2 detects *C. nebraskensis* strains, and; line C is a control line. The primer/probe sets for genus *Clavibacter* and *C. nebraskensis* are labeled with Biotin/DIG and Biotin/FAM, respectively. Step 1, both infected and healthy plant tissues were processed using TE buffer and a pestle; Step 2, the crude extract was directly used in RPA reactions as a template and incubated using closed hand palm for 30 minutes; Step 3, visual detection of multiplex RPA results on RPA-PCRD test strips displaying a specific detection of *Clavibacter* and *C. nebraskenis* from corn leaves in the upper PCRD, while Clavibacter detection from tomato stem in the lower PCRD. All the reactions were carried out using multiplex assays with TwistAmp-nfo kit. No cross-reactivity or negative results were observed despite using plant extract that contains plant inhibitors.

## DISCUSSION

In this study, we reported a reliable, specific, and sensitive multiplex RPA assay for the detection of the widely prevalent plant-pathogenic genus *Clavibacter*, in general, and more specifically, the re-emerged and increasingly important Goss’ Wilt pathogen, *C. nebraskensis.* With potential application for point-of-need detection in the field, this proof-of-concept assay based upon the combination of unique genomic regions and multiplex RPA detections coupled with a portable lateral flow device (LFD) may function as a potential tool to rapidly detect numerous targets without specialized equipment. Furthermore, in published literature on current RPA methods, there was no report of simultaneous detection of multiple plant pathogens in laboratory or field settings; thus a number of research studies may benefit from the findings of this study.

The ABC transporter ATP-binding protein CDS/ABC-transporter permease was selected for the detection of all known species and subspecies within the genus *Clavibacter*, and the MFS transporter gene for the specific detection of *C. nebraskensis*. These genomic regions were used previously by Dobhal et al, (2019) and Larrea-Sarmiento et. al (2019) to develop TaqMan qPCR and loop-mediated isothermal amplification-based assays and were proven to be unique, conserved, and highly specific. Besides previous genomic analyses, the regions have also been cross-checked with multiple genomes of each species and found to be specific (data not shown; 2, 33). Previously, RPA assays for plant bacterial pathogens using TwistAmp nfo chemistry coupled with LFD, have shown high specificity and ease of application (16, 20). The multiplex RPA assay developed here was tested with *Clavibacter*- and *Cn-*specific primer/probe sets using a total of 32 strains representing all known species/subspecies within the genus *Clavibacter*, including 14 strains of *C. nebraskensis*; the obtained results were specific (Table 1). The identity of these strains was previously confirmed using the *dnaA* gene region (2, 33).

Technical expertise along with instrument requirements and infrastructure limitations are common concerns when considering the practicality of diagnostic assays. RPA not only overcomes these limitations but also provides great advantages for on-site detection capability. The preclusion of DNA purification and the elimination of sophisticated equipment provides greater affordability to compensate for technology gaps and allows for the end-user to feasibly conduct multiplex detection tests in the field. TwistDx RPA reagents are available in lyophilized form and the kit was portable and user-friendly. Molecular analyses that use crude plant extracts as starting materials are often affected by inhibitory sample impurities (32). However, the remarkable tolerance to inhibitors offered by RPA led to the development of this rapid sample procedure for RPA assays utilizing tissue ground in TE buffer and applying the crude sap directly as a template (11, 16, 20).

The nature of the recombinase primer complexes used in RPA reactions enables amplification between 37-42 °C (13). The low incubation temperature and short reaction time (10 – 30 minutes) enhance the efficiency and flexibility of RPA for rapid multiplex pathogen detection (Figures 2 and 3). Although the intensity of the line on LFD was not intense, a positive reaction can be obtained in as little as 10 min. However, optimum amplification was observed when the RPA reaction was incubated for 20-30 min at 39 °C. Both Gamze et al, (2020) and Ahmed et al (2018) demonstrated optimum amplification at 30 min with no false positives or negatives. Previously in 2016, Arif and co-workers presented similar optimized test conditions with the *Rathayibacter toxicus*-specific RPA assay (14). The development of RPA based on equipment-free incubation using body temperature was employed previously by Crannell and collaborators for detection of intenstinal protozoa (2014) but this method has not been used for the detection of plant pathogens. This study showed successful, equipment-free incubation of RPA reactions at body temperature in the closed palm.

A thoroughly validated LAMP assay for specific detection of all known subspecies/species in the genus *Clavibacter* has already been developed by Dobhal and co-workers (2019), but the high-temperature requirement (ca. 65°C) limits the usefulness of LAMP without portable equipment designed for field applications; moreover, the LAMP assay was not able to discriminate *C. nebraskensis* from other *Clavibacter* species.

An optimized diagnostic tool must be sufficiently sensitive to detect the target within latently infected/ symptomless samples. The developed RPA assay detected the target sequences as low as 10 pg and 100 fg of genomic DNA of the genus *Clavibacter* (all species/subspecies) and *C. nebraskensis*, respectively. However, a 10-fold reduction in the sensitivity of the *Cn*-specific primer/probe set was observed when host sap was added into the 10-fold serially diluted genomic DNA (Figure 4). Previously, Ahmed et al (2018) reported an RPA assay for specific detection of genus *Pectobacterium* using TwistDx-nfo coupled with Milenia LFD; the assay showed detection as low as 10 fg from both sensitivity and spiked sensitivity assays. Detection limits of RPA assays are subject to distortion by background noise due to the formation of primer-dimers during the amplification of multiple targets or competition among the primers for the recombinase proteins (18, 27). Multiplex RPA assays were also optimized by adjusting the primer to probe ratio to maximize signal detection; equal concentrations of *Clavibacter-* and *Cn-* specific primer/probe sets displayed similar intensity lines across the strip, with no presence of false positives or false negatives.

Current multiplex RPA methodology is highly specific and sufficiently sensitive to detect multiple targets. The selection of unique targets by genome comparison resulted in specific primer/probe sets enabling the rapid and reliable performance of multiplex RPA. Additionally, efficient sample preparation generates results in about 40-50 minutes without DNA purification and sophisticated equipment. The developed multiplex RPA assay is ready to be used for point-of-need applications. The developed tools have potential applications in biosecurity, disease surveys, routine diagnostics, seed testing, and epidemiological studies.

## CONCLUSIONS

The multiplex RPA format coupled with LFD, which requires no nucleic acid purification or sophisticated instrumentation, can be performed at a low-temperature, with a short incubation time, and achieves sensitive detection makes the current multiplex RPA assay ideal for in-field and in-lab detection. Until now, multiplex RPA assays capable of accurate and simultaneous detection and differentiation of plant pathogens have not been published. The developed multiplex RPA assay has the potential for rapid and sensitive detection of multiple targets in field settings.

## ACKNOWLEDGMENTS

This work was supported by the USDA National Institute of Food and Agriculture, Hatch project 9038H, managed by the College of Tropical Agriculture and Human Resources. The strains used in this study were revived and further characterized with grant support from the National Science Foundation (NSF-CSBR grant no. DBI-1561663). The mention of trade names or commercial products in this publication does not imply recommendation or endorsement by the University of Hawaii. The granting agencies had no role in study design, data collection, and analysis, decision to publish, or preparation of the manuscript.

## REFERENCES

1. Yasuhara-Bell, J. and Alvarez, A.M. 2015. Seed-associated subspecies of the genus *Clavibacter* are clearly distinguishable from *Clavibacter michiganensis* subsp. michiganensis. Int J Syst Evol Microbiol 65, 811–826.

2. Dobhal, S., Larrea-Sarmiento, A., Alvarez, A. M., Arif, M. 2019. Development of a loop-mediated isothermal amplification assay for specific detection of all known subspecies of *Clavibacter michiganensis*. J Appl Microbiol; 126(2):388–401.

3. Li, X., Tambong, J., Yuan, K., Chen, W., Xu, H., Levesque, C.A. and De Boer, S.H. 2018. Re-classification of *Clavibacter michiganensis* subspecies on the basis of whole genome and multi-locus sequence analyses. Int J Syst Evol Microbiol 68, 234–240.

4. Tambong, J. T. 2017. Comparative genomics of *Clavibacter michiganensis* subspecies, pathogens of important agricultural crops. PLOS ONE, 12(3), e0172295.

5. Biddle, J., McGee, D., & Braun, E. 1990. Seed transmission of *Clavibacter michiganense* subsp. *nebraskense* in corn. Plant Disease, 74(11), 908–911.

6. Nemeth, J., Laszlo, E., & Emödy, L. 1991. *Clavibacter michiganensis* ssp. *insidiosus* in lucerne seeds. EPPO Bulletin, 21(4), 713–718.

7. Franc, G. 1999. Persistence and Latency of *Clavibacter michiganensis* subsp. *sepedonicus* in Field-Grown Seed Potatoes. Plant Disease, 83(3), 247–250.

8. Eichenlaub, R., & Gartemann, K. 2011. The *Clavibacter michiganensis* subspecies: Molecular investigation of Gram-positive bacterial plant pathogens. Annual Review of Phytopathology, 49(1), 445–464.

9. Crannell, Z., Castellanos-Gonzalez, A., Nair, G., Mejia, R., White, A. C., & Richards-Kortum, R. 2016. Multiplexed recombinase polymerase amplification assay to detect intestinal protozoa. Analytical Chemistry, 88(3), 1610–1616.

10. Wu, Y. D., Xu, M. J., Wang, Q. Q., Zhou, C. X., Wang, M., Zhu, X. Q., & Zhou, D. H. 2017. Recombinase polymerase amplification (RPA) combined with lateral flow (LF) strip for detection of Toxoplasma gondii in the environment. Veterinary Parasitology, 243, 199–203.

11. Lobato, I., & O’Sullivan, C. 2018. Recombinase polymerase amplification: Basics, applications and recent advances. TrAC Trends in Analytical Chemistry, 98, 19–35.

12. Notomi, T., Okayama, H., Masubuchi, H., Yonekawa, T., Watanabe, K., Amino, N., & Hase, T. 2000. Loop-mediated isothermal amplification of DNA. Nucleic acids research, 28(12), E63.

13. Piepenburg, O., Williams, C. H., Stemple, D. L., & Armes, N. A. 2006. DNA detection using recombination proteins. PLoS Biology, 4(7), e204.

14. Arif, M., Busot, G. Y., Mann, R., Rodoni, B., and Stack, J. P. 2016. Detection of the select agent *Rathayibacter toxicus* using recombinase polymerase amplification coupled with a lateral flow device. (Abstr.) Phytopathology 106:S4.23.

15. Song, J., Liu, C., Mauk, M. G., Rankin, S. C., Lok, J. B., Greenberg, R. M., & Bau, H. H. 2017. Two-stage isothermal enzymatic amplification for concurrent multiplex molecular detection. Clinical Chemistry, 63(3), 714–722.

16. Ahmed, F. A., Larrea-Sarmiento, A., Alvarez, A. M., & Arif, M. 2018. Genome-informed diagnostics for specific and rapid detection of *Pectobacterium* species using recombinase polymerase amplification coupled with a lateral flow device. Sci Rep 8, 15972.

17. Larrea-Sarmiento, A., Dhakal, U., Boluk, G., Fatdal, L., Alvarez, A., Strayer-Scherer, A., Paret, M., Jones, J. et al. (2018). Development of a genome-informed loop-mediated isothermal amplification assay for rapid and specific detection of *Xanthomonas euvesicatoria*. Sci Rep 8, 14298. doi.org/10.1038/s41598-018-32295-4.

18. Lau, H. Y., & Botella, J. R. (2017). Advanced DNA-Based Point-of-Care Diagnostic Methods for Plant Diseases Detection. Frontiers in Plant Science, 8, 2016.

19. Ocenar, J., Arizala, D, Boluk, G., Dhakal, U., Gunarathne, S., Paudel, S., Dobhal, S., & Arif, M. 2019. Development of a robust, field-deployable loop-mediated isothermal amplification (LAMP) assay for specific detection of potato pathogen *Dickeya dianthicola* targeting a unique genomic region. PLOS ONE 14(6): e0218868.

20. Boluk, G., Dobhal, S., Crockford, A. B., Melzer, M., Alvarez, A. M., & Arif, M. 2020. Genome-informed recombinase polymerase amplification assay coupled with a lateral flow device for in-field detection of *Dickeya* species. Plant Disease.

21. De Paz, H. D., Brotons, P., & Muñoz-Almagro, C. 2014. Molecular isothermal techniques for combating infectious diseases: towards low-cost point-of-care diagnostics. Expert Review of Molecular Diagnostics, 14(7), 827–843.

22. Clancy, E., Higgins, O., Forrest, M. S., Boo, T. W., Cormican, M., Barry, T., Piepenburg, O., & Smith, T. J. 2015. Development of a rapid recombinase polymerase amplification assay for the detection of *Streptococcus pneumoniae* in whole blood. BMC Infectious Diseases, 15, 481.

23. Daher, R. K., Stewart, G., Boissinot, M., & Bergeron, M. G. 2016. Recombinase polymerase amplification for diagnostic applications. Clinical Chemistry, 62(7), 947–958.

24. Kersting, S., Rausch, V., Bier, F. F., & von Nickisch-Rosenegk, M. 2014. Multiplex isothermal solid-phase recombinase polymerase amplification for the specific and fast DNA-based detection of three bacterial pathogens. Microchimica Acta, 181(13–14), 1715–1723.

25. Yang, Y., Qin, X., Sun, Y., Cong, G., Li, Y., & Zhang, Z. 2017. Development of isothermal recombinase polymerase amplification assay for rapid detection of *Porcine Circovirus* Type 2. BioMed Research International, 2017, 8403642.

26. Crannell, Z. A., Rohrman, B., & Richards-Kortum, R. 2014. Equipment-free incubation of recombinase polymerase amplification reactions using body heat. PLoS ONE, 9(11), e112146.

27. Kim, J. Y., & Lee, J.L. 2017. Development of a multiplex real-time recombinase polymerase amplification (RPA) assay for rapid quantitative detection of *Campylobacter coli* and jejuni from eggs and chicken products. Food Control, 73, 1247–1255.

28. Mekuria, T. A., Zhang, S., & Eastwell, K. C. 2014. Rapid and sensitive detection of Little cherry virus 2 using isothermal reverse transcription-recombinase polymerase amplification. Journal of Virological Methods, 205, 24–30.

29. Zhang, S., Ravelonandro, M., Russell, P., McOwen, N., Briard, P., Bohannon, S., & Vrient, 2014. Rapid diagnostic detection of plum pox virus in Prunus plants by isothermal AmplifyRP using reverse transcription-recombinase polymerase amplification. Journal of Virological Methods, 207, 114–120.

30. Miles, T. D., Martin, F. N., & Coffey, M. D. 2015. Development of Rapid Isothermal Amplification Assays for Detection of *Phytophthora* spp. in Plant Tissue, 105(2).

31. Londoño, M. A., Harmon, C. L., & Polston, J. E. 2016. Evaluation of recombinase polymerase amplification for detection of begomoviruses by plant diagnostic clinics. Virology Journal, 13(1), 48.

32. Kapoor, R., Srivastava, N., Kumar, S., Saritha, R. K., Sharma, S. K., Jain, R. K., & Baranwal, V. K. 2017. Development of a recombinase polymerase amplification assay for the diagnosis of banana bunchy top virus in different banana cultivars. Archives of Virology, 162(9), 2791–2796.

33. Larrea-Sarmiento, A., Alvarez, A. M., Stack, J. P. & Arif, M. 2019. Synergetic effect of non-complementary 5’ AT-rich sequences on the development of a multiplex TaqMan real-time PCR for specific and robust detection of *Clavibacter michiganensis* and *C*. *michiganensis* subsp. *nebraskensis*. PLOS ONE 14(7): e0218530.

34. Norman, D., & Alvarez, A. 1989. A rapid method for presumptive identification of *Xanthomonas campestris* pv. *dieffenbachiae* and other *Xanthomonas*. Plant Disease, 73(8), 654–658.

35. Altschul, S. F., Gish, W., Miller, W., Myers, E. W., & Lipman, D. J. 1990. Basic local alignment search tool. Journal of Molecular Biology, 215(3), 403–410.

36. Ahmad, A., Mbofung, G. Y., Acharya, J., Schmidt, C. L., Robertson, A. E. 2015. Characterization and comparison of *Clavibacter michiganensis* subsp. *nebraskensis* strains recovered from epiphytic and symptomatic infections of maize in Iowa. PLoS ONE 10(11): e0143553.

37. Kaneshiro, W. S., Mizumoto, C. Y., & Alvarez, A. M. 2006. Differentiation of *Clavibacter michiganensis* subsp. *michiganensis* from seed-borne saprophytes using ELISA, Biolog and 16S rDNA sequencing. Eur J Plant Pathol 116, 45–56.

