## Supplement Materials for "Multiplex recombinase polymerase amplification (RPA) assay developed using unique genomic regions and coupled with a lateral flow device for rapid on-site detection of genus *Clavibacter* and *C. nebraskensis*"

**Supplemental Figures**

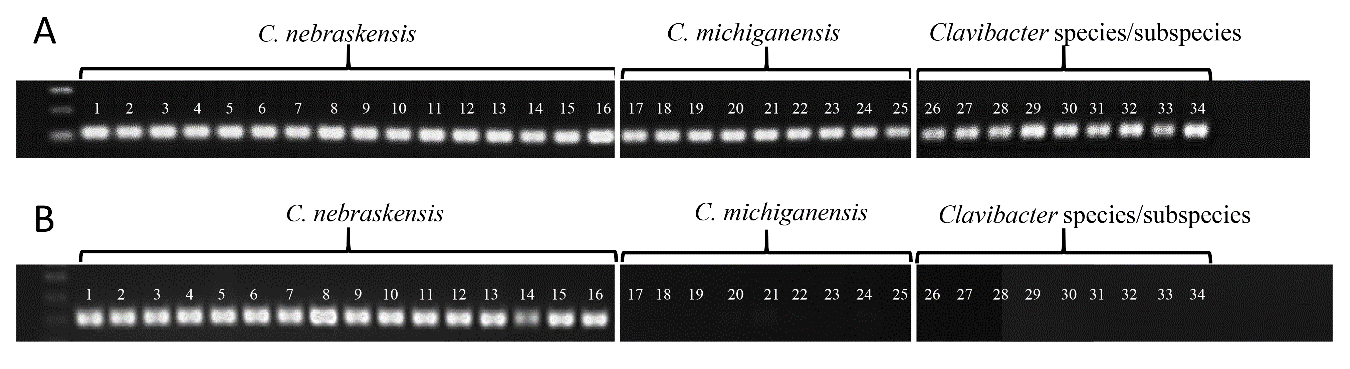

**Fig. S1.** Endpoint detection to evaluate the initial specificity of the developed primer sets. Thirty-four strains within the genus *Clavibacter* were used to verify the robustness of primer sets. A. *Clavibacter-*specific primers detected all 34 strains (2 DNA from infected samples). B. Specific detection of 16 strains of *C. nebraskensis* (samples 15 and 16 corresponded to DNA extracted from infected corn tissue). All strains and their characteristics are listed in Table 1. No amplification of the 20-non-*Cn* strains used in the exclusivity panel was observed with *Cn*-specific primers.

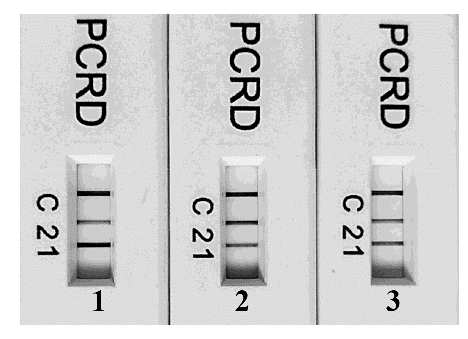

**Fig. S2.** RPA assay optimization for primer and probe concentration. Multiplex RPA reactions were performed following the provider instructions. *Clavibacter-*specific primer/probe concentrations were evaluated against *Cn*-specific primer/probe set. 1) low (*Cn* 1.2 µL/1.4 µL; *Clavibacter* 0.8 µL /0.6 µL); 2) equal (*Cn* 1.2 µL/0.8 µL; *Clavibacter* 1.2 µL/0.8 µL); and 3) high (*Cn* 1.5 µL/0.9 µL; *Clavibacter* 1.0 µL/0.6 µL).
